# Perfluorononanoic acid impedes mouse oocyte maturation by inducing mitochondrial dysfunction and oxidative stress

**DOI:** 10.1101/2021.07.06.451300

**Authors:** Xiaofei Jiao, Ning Liu, Yiding Xu, Huanyu Qiao

**Author notes:** These two authors contributed equally (Xiaofei Jiao, Ning Liu). Correspondence: Department of Comparative Biosciences, University of Illinois at Urbana-Champaign, 2001 South Lincoln Avenue, Urbana, IL 61802, USA.

## Abstract

Perfluorononanoic acid (PFNA), a member of PFAS, is frequently detected in human blood and tissues, even in follicular fluid of women. The exposure of PFNA, but not PFOA and PFOS, is positively correlated with miscarriage and increased time to pregnancy. Toxicological studies indicated that PFNA exposure is associated with immunotoxicity, hepatotoxicity, developmental toxicity, and reproductive toxicity in animals. However, there is little information regarding the toxic effects of PFNA on oocyte maturation. In this study, we investigated the toxic effects of PFNA exposure on mouse oocyte maturation *in vitro*. Our results showed that 600 μM PFNA significantly inhibited germinal vesicle breakdown (GVBD) and polar body extrusion (PBE) in mouse oocytes. Our further study revealed that PFNA induced abnormal metaphase I (MI) spindle assembly, evidenced by malformed spindles and mislocalization of p-ERK1/2 in PFNA-treated oocytes. We also found that PFNA induced abnormal mitochondrial distribution and increased mitochondrial membrane potential. Consequently, PFNA increased reactive oxygen species (ROS) levels, leading to oxidative stress, DNA damage, and eventually early-stage apoptosis in oocytes. In addition, after 14 h culture, PFNA disrupted the formation of metaphase II (MII) spindle in most PFNA-treated oocytes with polar bodies. Collectively, our results indicate that PFNA interferes with oocyte maturation *in vitro* via disrupting spindle assembly, damaging mitochondrial functions, and inducing oxidative stress, DNA damage, and early-stage apoptosis.

## 1. Introduction

Per- and polyfluoroalkyl substances (PFAS) are ubiquitous and negatively affect the female reproductive system. Perfluoroalkyl acids (PFAAs), a subgroup of PFAS, are a family of man-made industrial compounds that have been widely used in consumer products, including non-stick cookware, fire-retardant foams, water- and stain-resistant coating, and food packaging materials due to their anti-wetting and surfactant properties [1]. Humans are exposed to PFAAs from different sources, such as contaminated air, dust, food, and drinking water [2–4]. Lines of evidence suggest that PFAA exposure is associated with adverse health effects including carcinogenesis, hepatotoxicity, immunotoxicity, neurotoxicity, endocrine disruption, developmental toxicity, and reproductive toxicity [5–7]. Most PFAAs can accumulate in human tissues and are resistant to metabolic degradation [8]. Because of their persistence and toxic effects, many PFAAs are considered as persistent organic pollutants – “forever chemicals”. Thus, two major types of PFAAs, perfluorooctane sulfonate (PFOS) and perfluorooctanoic acid (PFOA), were phased out of production in the USA. However, hundreds of other PFAAs are still produced and used without restrictions, partially due to a lack of toxicity studies.

Perfluorononanoic acid (PFNA) is a long-chain PFAA containing a nine-carbon backbone. PFNA, like PFOA and PFOS, is an environmental contaminant that can be detected in animals and humans [9,10], even in human follicular fluid (0.2-2.1 ng/mL)[11]. Although the detectable level of PFNA (average 1.49 ng/mL in the US) is much lower than those of PFOA and PFOS in human blood, its levels have been steadily increasing from 1999 to 2004 [12,13]. From 2003 to 2014, there was a slight decrease of PFNA levels in NHANES, but the trend was not significant [14]. Even in wild animals, PFNA levels have increased over time [15,16]. Long-chain PFAAs, such as PFNA, are suggested to have higher toxicities compared to their short-chain counterparts [17,18]. Thus, PFNA has attracted wide public attention in recent years.

Toxicological studies have indicated that PFNA exposure is associated with many adverse effects. Exposure to PFNA resulted in toxic effects on lymphoid organs and lymphocyte homeostasis through activating the peroxisome proliferator-activated receptors (PPARs) and hypothalamic-pituitary-adrenal (HPA) axis in BALB/c mice [19]. Administration of PFNA to BALB/c mice also led to abnormal liver enlargement and disturbance of hepatic lipid [20]. Das et al. [21] showed that fetal exposure of PFNA during pregnancy postponed the onset of puberty, increased liver weight, and lowered survival rates among F1 offspring of CD-1 mice. Recent studies showed that chronic and acute PFNA treatment impaired spermatogenesis and sperm quality by disrupting testosterone biosynthesis and inducing oxidative stress in male mice [22,23]. In addition, PFNA may cause cytotoxicity in JEG-3 cells, PC12 cells, and lung cells *in vitro* [18,24,25]. Furthermore, PFNA causes developmental toxicity by impairing bovine embryo growth *in vitro* [26]. All the aforementioned studies suggest that PFNA exerts toxic effects both *in vivo* and *in vitro*. However, whether PFNA can affect oocyte maturation remains unknown.

In this study, we used a mouse oocyte *in-vitro* culture system to study the toxic effects of PFNA exposure on oocyte maturation. The germinal vesicle breakdown (GVBD) rate, polar body extrusion (PBE) rate, spindle assembly, chromosome alignment, ROS levels, and DNA damage were examined to characterize the adverse effects mediated by PFNA.

## 2. Materials and methods

### 2.1 Chemicals and antibodies

Perfluorononanoic acid (PFNA, 97%), 3-Isobutyl-1-methylxanthine (IBMX, ≥ 99%) and 2’,7’-dichlorofluorescin diacetate (DCFH-DA, ≥ 97%) were obtained from Sigma-Aldrich (St. Louis, MO, USA). JC-1 Mitochondrial Membrane Potential Detection Kit and MitoView™ Green were purchased from Biotium (Hayward, CA, USA). Mouse monoclonal anti-α-Tubulin-FITC antibody and anti-phospho-Histone H2A.X (Ser139) antibody, clone JBW301 were purchased from Sigma-Aldrich (St. Louis, MO, USA). Anti-phospho-p44/42 MAPK (Erk1/2) antibody was purchased from Cell Signaling Technologies. Alexa Fluor 488-conjugated goat anti-mouse IgG (H + L) and Alexa Fluor 594-conjugated goat anti-rabbit IgG (H + L) secondary antibodies were obtained from Molecular Probes.

### 2.2 Animals

CD-1 mice (Charles River Laboratories, Wilmington, MA) were used in this study and housed in the Animal Care Facility at the University of Illinois Urbana-Champaign (UIUC). Mice were housed under 12 h dark/12 h light cycles at 22 ± 1°C and were provided food and water ad libitum. Animal handling and procedures were approved by the UIUC Institutional Animal Care and Use Committee.

### 2.3 Mouse oocyte *in vitro* culture

For *in vitro* maturation (IVM), 3-4-week-old female mice were euthanized, and ovaries were isolated using a dissection microscope. After washing the ovaries three times with pre-warmed M2 (Sigma-Aldrich, St. Louis, MO, USA) medium containing IBMX (100 μM), the cumulus-oocyte complexes (COCs) were isolated by rupturing the antral ovarian follicles with sterile needles. To obtain denuded germinal vesicle (GV) intact oocytes, cumulus cells were removed by repeated pipetting. Fully-grown oocytes were collected in pre-warmed M2 supplemented with 100 μM IBMX. After three washes in the culture medium (M16), oocytes were transferred into a small drop of M16 medium (Sigma-Aldrich, St. Louis, MO, USA) under mineral oil and cultured at 37°C in a 5% CO2 incubator. After being cultured for 2 h, 8 h, and 14 h, most oocytes reach germinal vesicle breakdown (GVBD), metaphase I (MI), and metaphase II (MII), respectively. Next, oocytes at different stages were selected for data analysis according to the experimental design.

### 2.4 Chemical treatment

The lowest concentration (3 nM) of PFNA used in this study is the environmentally relevant dose that detected in human follicular fluid [11]. The highest concentration (600 μM) of PFNA was based on our experimental results (Fig. 1B and C) and published research [18]. PFNA powder was dissolved in DMSO and diluted into M16 medium to final concentrations of 3 nM, 200 μM, 400 μM and 600 μM. The control medium was supplemented with the same amount of DMSO and the final concentration of the solvent was no more than 0.1% in all of the culture media.

**Figure 1.**
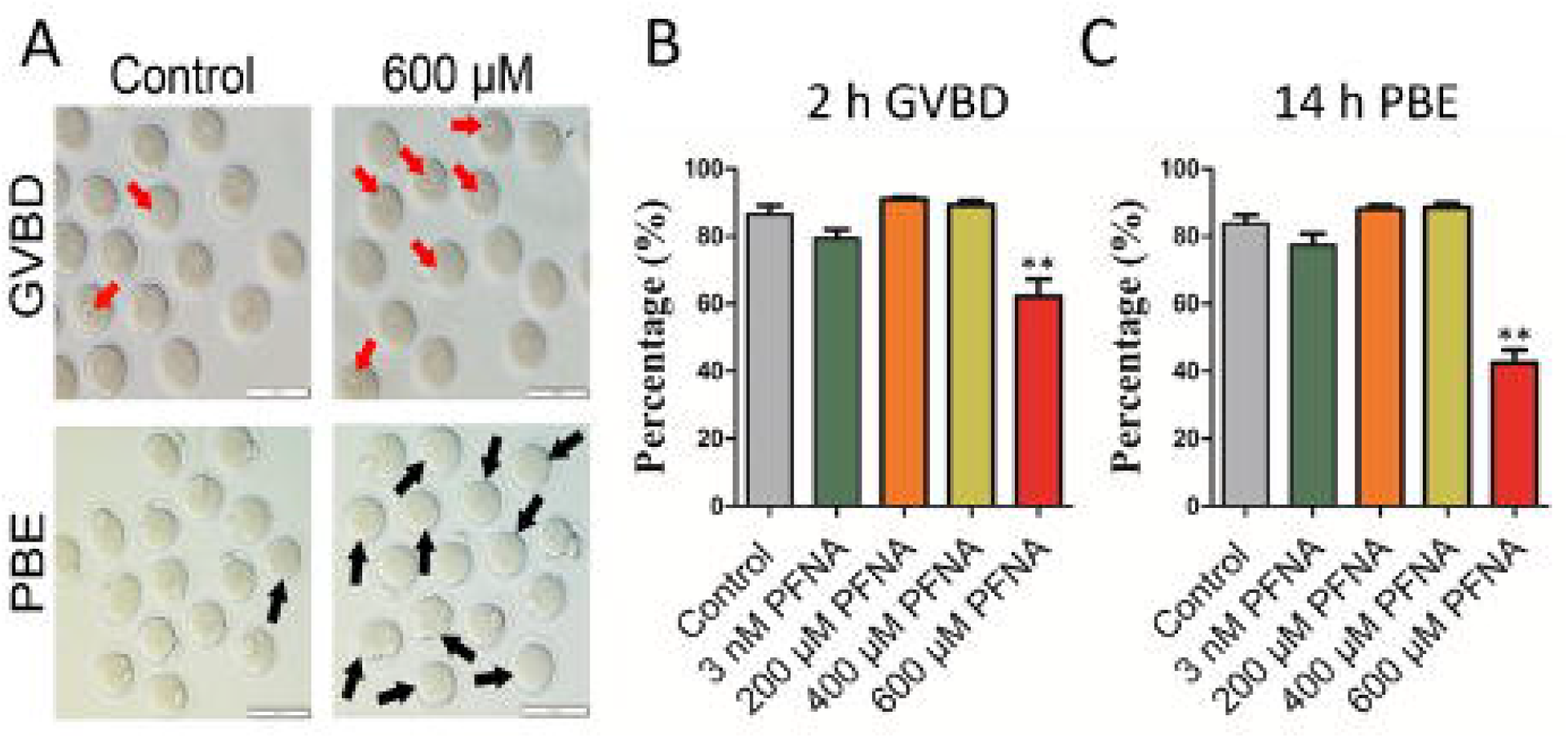
PFNA inhibited meiotic progression in mouse oocytes. (A) Representative images showed the morphology of oocytes after 2 h and 14 h *in vitro* culture in the control and PFNA-treated groups. The red arrows highlighted the oocytes with germinal vesicle. The black arrows pointed out the oocytes that failed to extrude their first polar bodies. Scale bar, 100 μm. (B) The rates of GVBD in the untreated control and PFNA-treated groups (3 nM, 200 μM, 400 μM, 600 μM) after 2-hour culture. (C) The PBE rates in the untreated control and PFNA-treated groups (3 nM, 200 μM, 400 μM, 600 μM) after 14-hour culture. A total of 180 oocytes in the control, 85 oocytes in the 3 nM group, 111 oocytes in the 200 μM group, 108 oocytes in the 400 μM group, and 196 oocytes in the 600 μM group were analyzed for GVBD and PBE rates. Data were presented as mean ± SEM of at least three independent experiments. \*\*\**P* < 0.001, compared with control.

### 2.5 Immunofluorescent staining

Oocytes were fixed in a large drop of 4% (W/V) paraformaldehyde in phosphate buffered saline (PBS) for 30 min at room temperature. After fixation, the oocytes were washed with washing buffer (0.1% tween-20 and 0.01% Triton X-100 in PBS) three times, then permeabilized with 0.5% Triton X-100 in PBS for 20 min at room temperature. After being washed twice, the oocytes were transferred into a drop of blocking buffer containing 3% bovine serum albumin (BSA) for one hour at room temperature. Then, the oocytes were washed three times and incubated with FITC-labelled mouse anti-α-tubulin (1:200, Sigma-Aldrich) for 1 h at 37°C.

For γ-H2AX staining, oocytes were incubated overnight at 4°C with mouse monoclonal anti-γ-H2AX antibody, clone JBW301 (1:200, Millipore-Sigma). After three times (10 min each) of washing, the oocytes were incubated with the FITC-conjugated goat anti-mouse IgG (H + L) secondary antibodies (1:300, Molecular Probes) for 1 h at 37°C. The oocytes were again washed three times (10 min each) after the incubation at 37°C, then oocyte DNA was counter-stained with 1 μg/ml of 4’,6-diamidine-2’-phenylindole dihydrochloride (DAPI) for 10 min at room temperature. After being washed twice with washing buffer, the oocytes were mounted on glass slides with 80% glycerol and examined with a Nikon A1R confocal microscope. The images were processed by using NIS-Elements software.

### 2.6 Measurement of intracellular reactive oxygen species (ROS) levels

An oxidation-sensitive fluorescent probe, 2’,7’-dichlorofluorescin diacetate (DCFH-DA) was used to determine the intracellular ROS level in both control and PFNA-treated oocytes. Briefly, oocytes were transferred into M2 medium containing 5 μM DCFH-DA and incubated at 37°C in a 5% CO2 incubator for 30 min. After incubation, the oocytes were washed twice in fresh M2 medium to remove the DCFH-DA, the signal was examined immediately by a fluorescence microscope (Olympus, IX73), and the images of the control and PFNA-treated oocytes were taken using the same imaging parameters. To quantify the signal intensity, images were processed by using the Image J software (National Institutes of Health, USA) to indicate the intensity of the region of interest (ROI) in the images.

### 2.7 Measurement of mitochondrial membrane potential (MMP)

The JC-1 mitochondrial membrane potential detection kit (Biotium, Hayward, CA, USA) was used for measuring mitochondrial membrane potential in oocytes. Oocytes were washed in fresh M2 medium 2-3 times and then incubated in M2 medium containing 1X JC-1 solution at 37°C for 30 min in the dark. Then the oocytes were washed again 2-3 times in fresh M2 medium and the signal was examined immediately by a fluorescence microscope (Olympus, IX73). Images of the control and PFNA-treated oocytes were taken using the same imaging parameters. For the quantification of the signal intensity, images were processed by using the Image J software (National Institutes of Health, USA) to indicate the intensity of the ROI in the images.

### 2.8 Mitochondrial staining

MitoView^™^ Green (Biotium, Hayward, CA, USA) was used to stain mitochondria in oocytes. Briefly, oocytes were washed in fresh M2 medium 2-3 times and then incubated in M2 medium containing 200 nM MitoView^™^ Green at 37°C for 30 min in the dark. Then oocytes were washed 2-3 times in fresh M2 medium. Next, the green signals from the control and PFNA-treated oocytes were immediately examined by a Nikon A1R confocal microscope using the same imaging parameters, and then processed using NIS-Elements software.

### 2.9 Detecting oocyte death by annexin-V staining

As described by Zhang et al., (2019), we used Annexin-V-FITC Apoptosis Detection Kit (eBioscience, San Diego, CA, USA) to identify apoptotic oocytes. After being cultured for 14 hours with or without PFNA, mouse oocytes were washed with fresh M2 medium and then incubated in the binding buffer for 5 min at room temperature. Next, the oocytes were incubated in annexin-V solution (1:40 in the binding buffer) at room temperature for 20 min in dark. Then oocytes were washed with M2 medium and incubated in the binding buffer for 5 min to remove unbound annexin-V. Without oocyte fixation, fluorescent images of oocytes were taken by using the same imaging parameters under Nikon A1R Confocal Microscope. To quantify the fluorescent intensity of annexin-V in the treated and control oocytes, images were processed by using the NIS-Elements software to measure the mean annexin-V signal intensity within zona pellucida of each oocyte. The relative fluorescent intensity was calculated by dividing all data points by the average intensity of the control group in each experiment.

### 2.10 Statistical analysis

Each treatment was conducted with at least three replicates. The data was presented as mean ± SEM. Comparisons between multiple groups were analyzed by one-way analysis of variance (ANOVA), followed by Tukey post comparison, and unpaired two-tailed t-test was applied to analyze the data between two different groups using Graph-Pad Prism analysis software. Comparisons were considered significant at \**P* < 0.05, \*\**P* < 0.01 and \*\*\**P* < 0.001. All charts were plotted by Graph-Pad Prism 5.0 (San Diego, USA).

## 3 Results

### 3.1 PFNA inhibited meiotic progression in mouse oocytes

To examine the effects of PFNA exposure on oocyte maturation *in vitro*, a range of concentrations of PFNA (0 μM, 3 nM, 200 μM, 400 μM and 600 μM) were used to treat mouse oocytes. The rates of germinal vesicle breakdown (GVBD) and polar body extrusion (PBE) were recorded at 2 h and 14 h of the oocyte culture, respectively. Our results showed that the 3 nM (environmentally relevant concentration), 200 μM and 400 μM PFNA treatments had no significant effects on GVBD rate compared to the control (*P* > 0.05) (Fig. 1). However, the 600 μM PFNA significantly reduced the GVBD rate in the exposed oocytes compared to the control after 2 h *in-vitro* culture (70.75 ± 3.67% in the 600 μM PFNA versus 91.37 ± 4.38% in the control, *P* < 0.001) (Fig. 1A & B).

Next, we calculated the proportion of oocytes with a polar body to get the PBE rates of the different treatment groups. Our results showed that most of the oocytes treated with the 3 nM PFNA, 200 μM PFNA, 400 μM PFNA, and vehicle control, extruded their polar bodies normally (*P* > 0.05). However, the 600 μM PFNA treatment severely inhibited polar-body extrusion in exposed oocytes, which was evidenced by a much lower PBE rate compared with control (33.95 ± 2.88% in the 600 μM PFNA versus 88.35 ± 2.42% in the control, *P* < 0.001) (Fig. 1A & C). According to the above results, we selected the 600μM concentration of PFNA for subsequent analysis because there was a significant decrease on the GVBD and PBE rates at this concentration.

### 3.2 PFNA induced spindle abnormality, clustering defects of microtubule organizing center (MTOC), and chromosome misalignment in MI mouse oocytes

Since there was a significant reduction of PBE rate in PFNA-treated oocytes and spindle assembly is essential for polar-body extrusion, we next examined whether abnormal spindle assembly at MI stage results in PBE failure in treated oocytes. Our results showed that the control oocytes displayed typical barrel-shaped spindles, but many PFNA-treated oocytes contained malformed spindles (examples of malformed spindles, such as elongated spindles, have been shown in Fig. 2A and 2D). Statistically, the abnormal spindle rate in the PFNA-treated group was significantly higher than that in the control (43.31 ± 3.55% in the 600 μM PFNA group versus 14.95 ± 2.80% in the control, *P* < 0.001; Fig. 2B). In addition, abnormal spindle assembly in the PFNA-exposed oocytes was identified by the mislocalization of p-ERK1/2, MTOC-associated proteins that are enriched in spindle poles to regulate proper spindle formation during meiosis [28]. As shown in Fig. 2D, p-ERK1/2 was only localized at the two poles of the spindle in control oocytes. In contrast, several additional p-ERK1/2 locations away from the two poles in the PFNA-treated oocytes were highlighted by white arrows in Fig. 2D, indicating MTOC clustering defects. Consequently, chromosome alignment was also disrupted by PFNA exposure. Our results showed that the chromosomes were well aligned at the MI plate in the control group. However, in the PFNA-exposed oocytes, chromosomes were not tightly aligned at the MI plate (see the arrows) (Fig. 2A). Statistical analysis also revealed that the rate of abnormal chromosome alignment in the PFNA-treated group was significantly higher than that in the control group (29.92 ± 10.65% in the 600 μM PFNA group versus 9.70 ± 3.37% in the control, *P* < 0.05; Fig. 2C).

**Figure 2.**
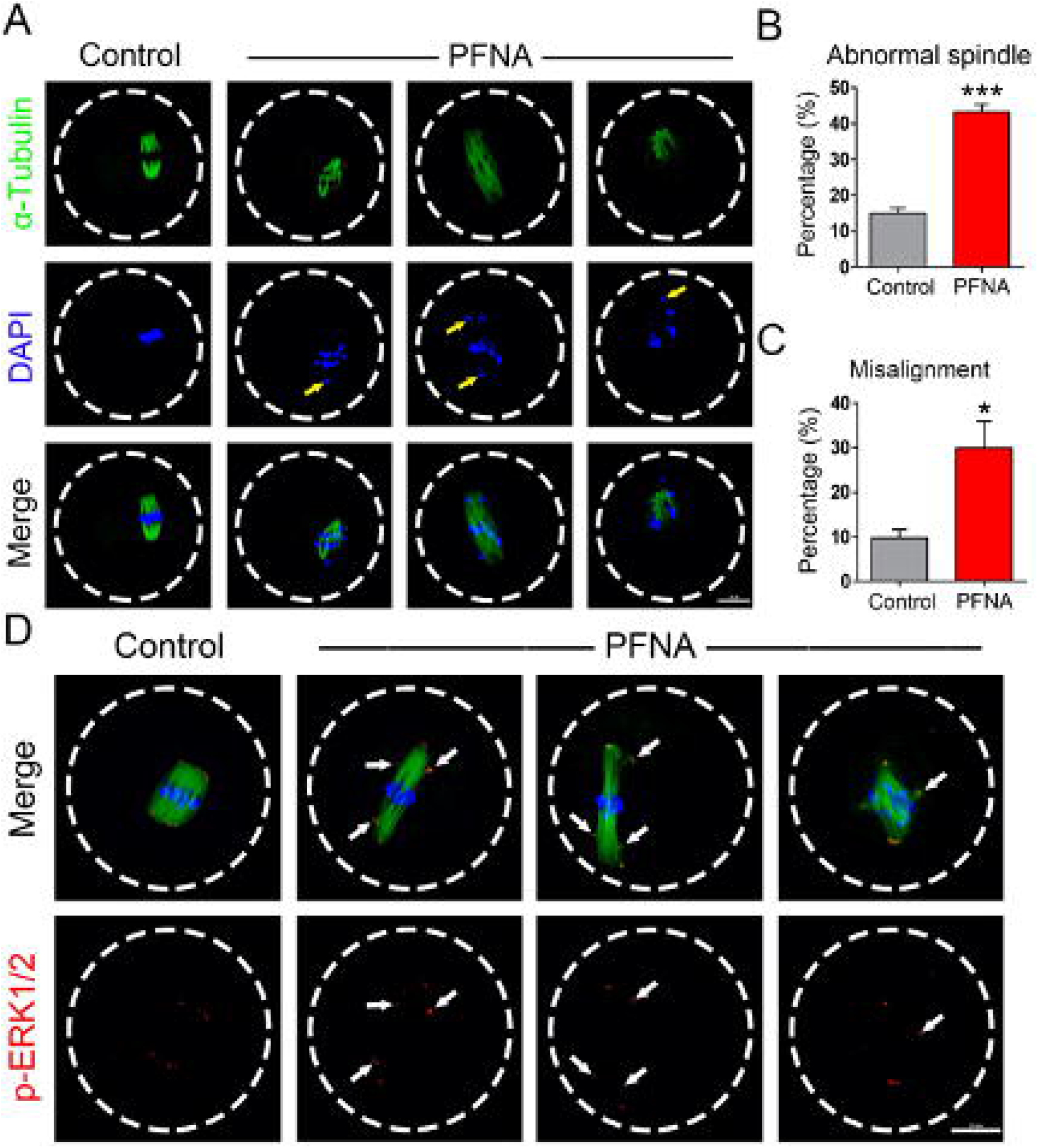
PFNA induced spindle abnormality, MTOC clustering defects, and chromosome misalignment in MI mouse oocytes. (A) Representative images showed the morphology of spindles and chromosomes after 8 h *in vitro* culture in the control and 600 μM PFNA-treated groups. The yellow arrows highlighted the misaligned chromosomes. Scale bar, 20 μm. (B) The rates of abnormal spindle in the untreated control and 600 μM PFNA-treated groups. (C) The rates of chromosome misalignment in the control and 600 μM PFNA-treated groups. (D) Images showed the p-ERK1/2 localization in the control and 600 μM PFNA-treated oocytes. The white arrows indicated the mislocalization of p-ERK1/2; α-Tubulin, green; p-ERK1/2, red; chromosomes/DNA, blue. Scale bar, 20 μm. A total of 41 oocytes in the control and 46 oocytes in the 600 μM PFNA-treated groups were analyzed for the abnormal spindle and chromosome misalignment rates. Data were presented as mean ± SEM of at least three independent experiments. t-test, \**P* < 0.05 and \*\*\**P* < 0.001, compared with control.

### 3.3 PFNA treatment resulted in mitochondrial dysfunction in mouse oocytes

Previous studies indicated that PFNA exposure damaged mitochondria in lung cells [18]. Thus, we hypothesized that PFNA affect oocyte maturation by impairing mitochondrial functions. To test our hypothesis, we used Mito-view dye, a mitochondrial stain, to check mitochondrial distribution. Our results showed that the mitochondria were evenly distributed as small dots in the untreated control oocytes, whereas most of the mitochondria aggregated in certain sites in the PFNA-treated oocytes (Fig. 3A). The proportion of oocytes with abnormal mitochondrial distribution in the treated oocytes was significantly higher than that in the control oocytes (71.38 ± 14.72% in the 600 μM PFNA group versus 13.33 ± 5.77% in the control, *P* < 0.01; Fig. 3B), suggesting that PFNA disrupted mitochondrial distribution in oocytes.

**Figure 3.**
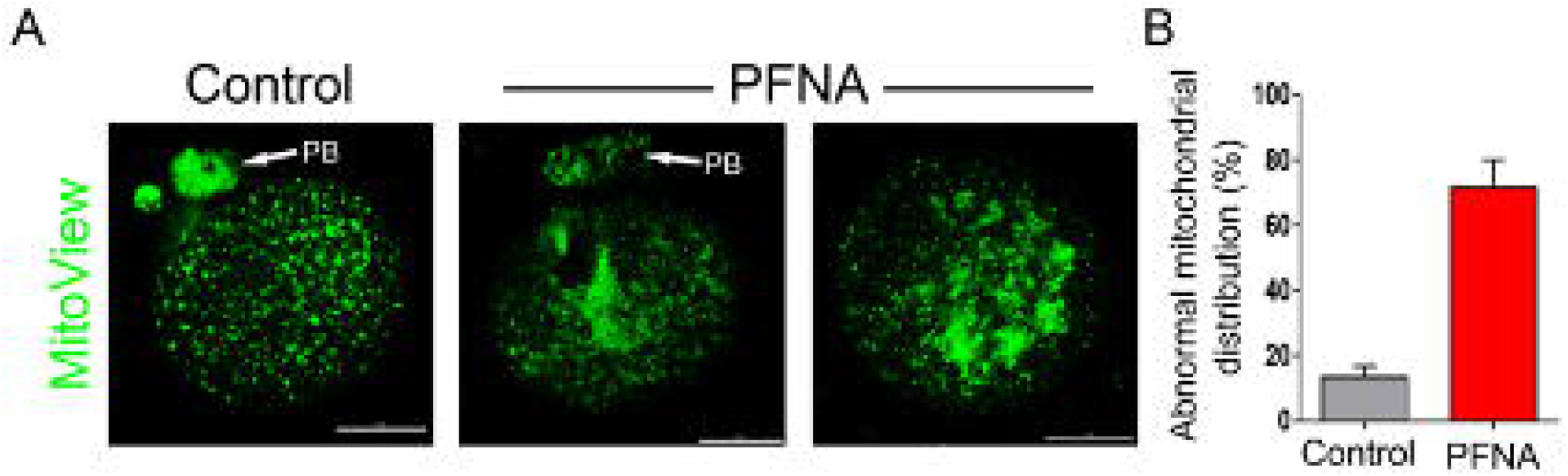
PFNA induced mitochondrial dysfunction in mouse oocytes. (A) Representative images showed the mitochondrial distribution patterns in control and 600 μM PFNA-exposed oocytes. A total of 30 oocytes in the control and 31 oocytes in the 600 μM PFNA-treated groups were analyzed for mitochondrial distribution analysis. PB, polar body; MitoView (a fluorogenic mitochondrial dye), green. Scale bar, 20 μm. (B) The proportions of oocytes with abnormal mitochondrial distribution patterns in the control and the 600 μM PFNA-treated groups.

### 3.4 PFNA increased reactive oxygen species (ROS) levels and damaged DNA in mouse oocytes

The biochemical reactions within mitochondria are major resources to generate endogenous ROS in cells. To further study mitochondrial functions and its association with oxidative stress, we examined mitochondrial membrane potential, a key factor of mitochondrial function, in both the control and PFNA-treated MI oocytes by using JC-1 staining. JC-1 dye is lipophilic and naturally exhibits green fluorescence. JC-1 starts to form reversible complexes (J aggregates) when JC-1 accumulates in mitochondria and reaches a certain concentration. The J aggregates exhibit red fluorescence. The red/green fluorescence ratio of JC-1 dye in the mitochondria can be considered as a direct assessment of the mitochondria polarization state. The stronger the red fluorescent signal of JC-1 the higher the mitochondrial membrane potential. Our results showed that the red signals of JC-1 staining in PFNA-exposed oocytes were brighter than that in the control oocytes (Supplemental Fig. 1A). Statistically, the ratio of red signal/green signal was significantly higher in the treated oocytes than that in the control (2.37 ± 0.73 in the 600 μM PFNA group versus 1.33 ± 0.25 in the control, *P* < 0.001; Supplemental Fig. 1B), indicating that PFNA elevated mitochondrial potential in oocytes.

High mitochondrial membrane potential is positively correlated with ROS overgeneration that leads to oxidative stress [29]. Thus, we hypothesized that PFNA treatment leads to oxidative stress in the PFNA-treated oocytes. To test this hypothesis, we checked the ROS levels in the control and PFNA-exposed groups by using DCFH-DA, a specific probe for ROS detection. Our results showed that the signal of DCFH-DA in the PFNA-exposed oocytes was much brighter than that in the control oocytes (Fig. 4A). Statistical analysis revealed that the intensity of DCFH-DA in the PFNA-treated oocytes was significantly higher than that in the control (Fig. 4B), which indicated that PFNA elevated ROS levels in the treated oocytes.

**Figure 4.**
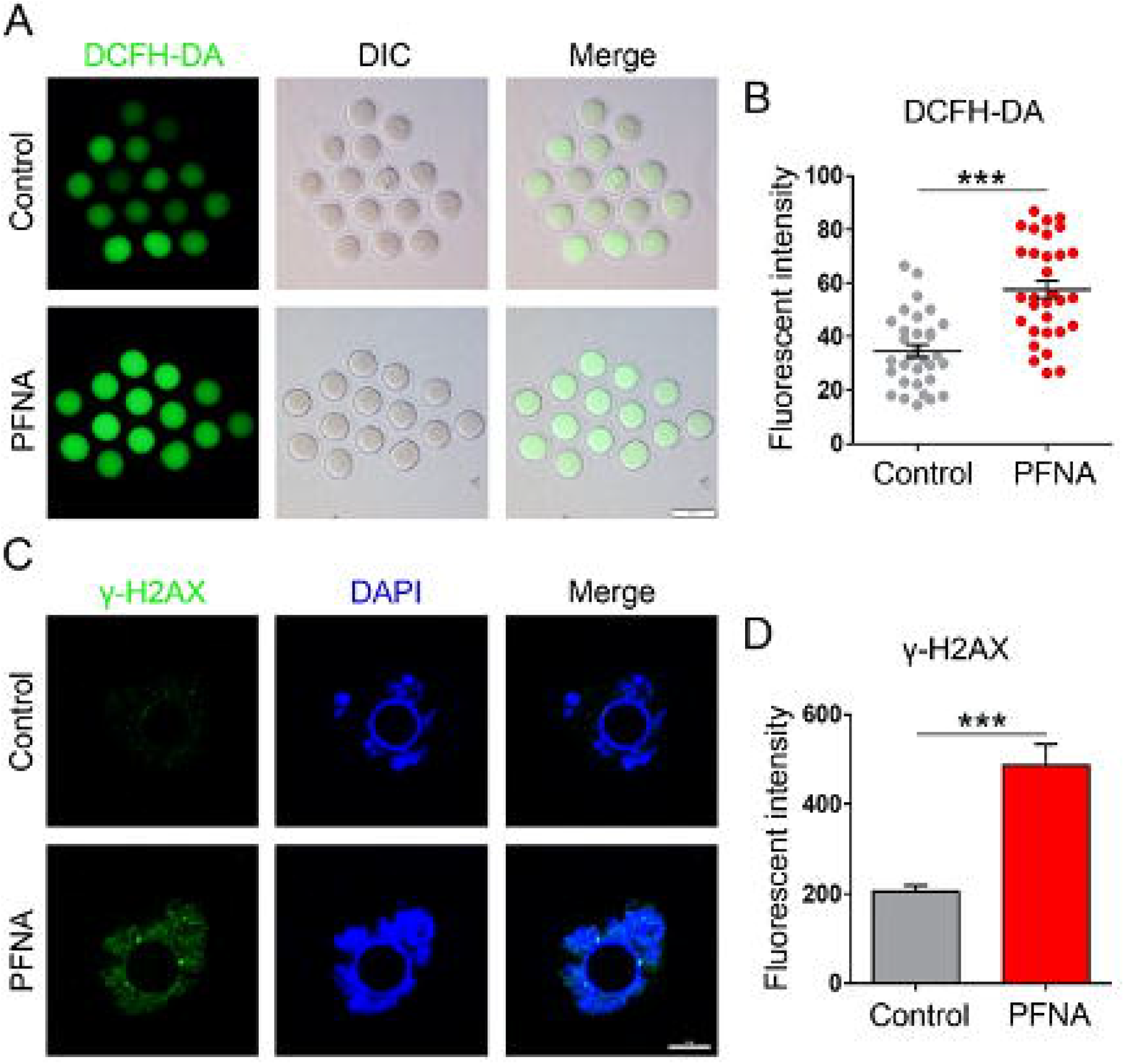
PFNA treatment increased ROS level and induced DNA damage in mouse oocytes. (A) Representative images of DCHF-DA fluorescence (green) in the control and 600 μM PFNA-exposed oocytes. DIC, differential interference contrast. Scale bar, 100 μm. (B) Quantitative analysis of DCHF-DA fluorescence intensity in the control and 600 μM PFNA-treated groups. A total of 32 oocytes in the control and 31 oocytes in the 600 μM PFNA-treated group were used for ROS level analysis. (C) Representative images showed γ-H2AX signal in the control and 600 μM PFNA-treated oocytes. Green, γ-H2AX; blue, DNA. Scale bar = 10 μm. (D) statistical analysis of average green intensity in the control and PFNA-exposed oocytes. A total of 29 oocytes in the control and 34 oocytes in the 600 μM PFNA-treated group were used for γ-H2AX signal intensity analysis. Data were presented as mean percentage (mean ± SEM) of at least three independent experiments. t-test, \*\*\**P* < 0.001, compared with control.

Excessive ROS in oocytes can induce DNA damage. Thus, we checked whether PFNA finally resulted in DNA damage in oocytes. We used IBMX to arrest oocytes at the GV stage and cultured them with and without PFNA for 8 h, then the arrested oocytes were collected for γ-H2AX staining, a marker for DNA double-stranded breaks. Our results showed that the green (γ-H2AX) signal was much brighter in the PFNA-exposed oocytes than that in the control (Fig. 4C). Statistically, the average γ-H2AX signal intensity in the PFNA-exposed oocytes was significantly higher than that in the control (Fig. 4D). Altogether, the data suggested that PFNA exposure led to DNA damage in mouse oocytes.

### 3.5 PFNA treatment caused metaphase II spindle abnormality in mouse oocytes

Some PFNA-treated oocytes still could extrude their polar bodies. Therefore, we checked the quality of these oocytes with polar bodies. The oocytes with polar bodies in both control and PFNA-treated groups were stained with α-tubulin-FITC antibody and DAPI to analyze the spindle morphology. Spindles in the control oocytes displayed a typical barrel shape; whereas spindles were disorganized and even totally disappeared in the PFNA-exposed group (Fig. 5A). Statistically, the abnormal spindle rate was significantly higher in the PFNA-treated group than in the control (77.81 ± 6.94% in the 600 μM PFNA group versus 16.27 ± 8.94% in the control, *P* < 0.001; Fig. 5B). In addition, chromosome misalignment was observed in the PFNA-treated oocytes, indicated by the arrows (Fig. 5A). Statistically, the rate of abnormal chromosome alignment in the PFNA-treated group was significantly higher than that in the control group (59.31 ± 8.07% in the 600 μM PFNA group versus 16.27 ± 8.94% in the control, *P* < 0.001; Fig. 5C).

**Figure 5.**
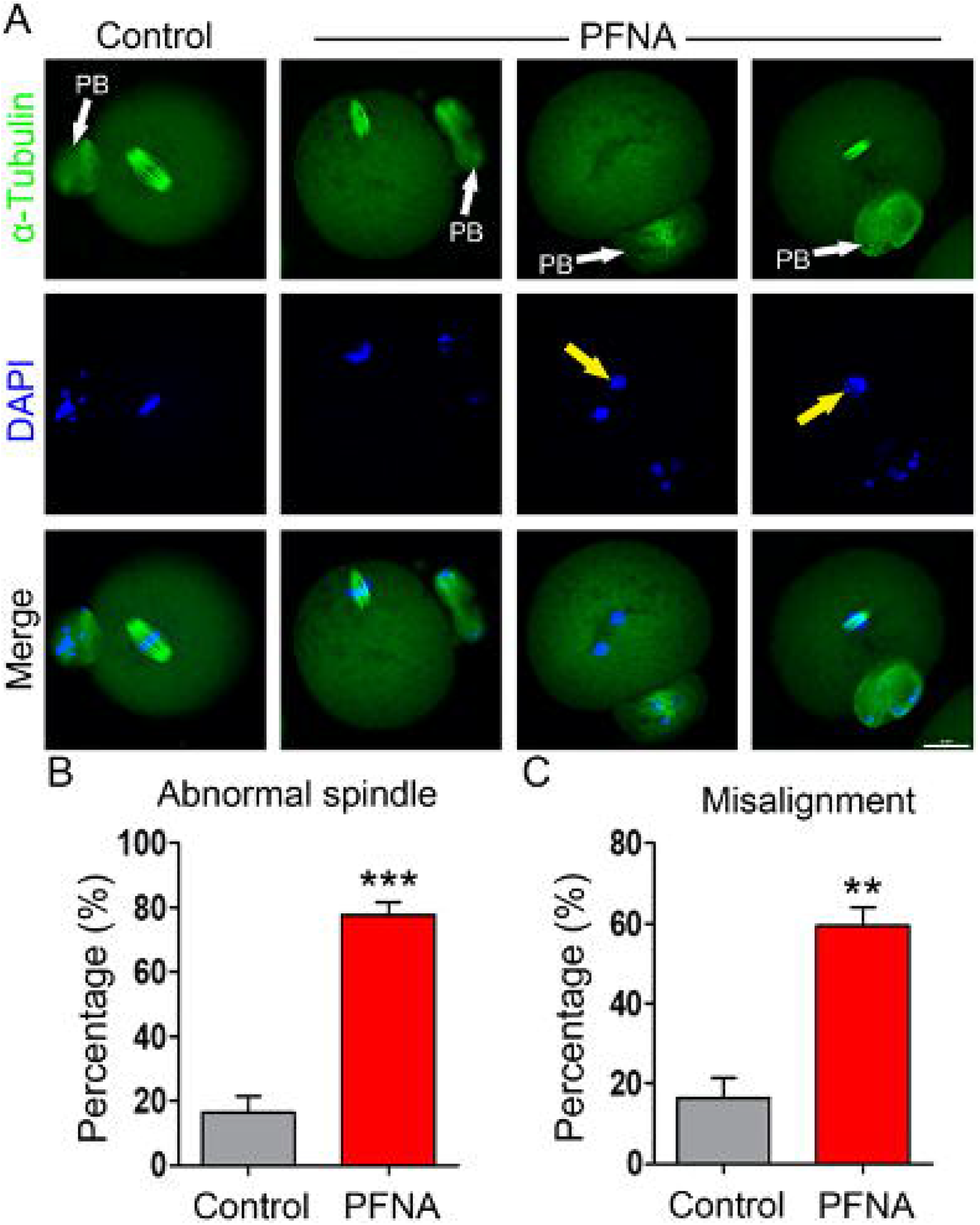
PFNA disrupted metaphase II spindle in mouse oocytes. (A) Representative images showed the spindle morphology and chromosome alignment in the untreated control and 600 μM PFNA-exposed oocytes. PB, polar body; α-tubulin, green; chromosomes/DNA, blue. Scale bar, 20 μm. Yellow arrows highlight the misaligned chromosomes. (B) The rates of abnormal MII spindle assembly in the control and 600 μM PFNA-treated groups. (C) The rates of chromosome misalignment in the control and 600 μM PFNA-treated groups. A total of 34 oocytes in the control and 33 in the 600 μM PFNA-treated groups were used for analysis. Data were presented as mean percentage (mean ± SEM) of at least three independent experiments. t-test, \*\**P* < 0.01 and \*\*\**P* < 0.001, compared with control.

### 3.6 PFNA treatment induced early-stage apoptosis in mouse oocytes

It has been reported that elevated ROS levels could promote oocyte apoptosis [30,31]. In addition, DNA damage and chromosome mis-segregation could also result in apoptosis [32,33]. Thus, we hypothesized that the oxidative stress, DNA damage, and chromosome misalignment induced by PFNA could cause oocyte death. To test this hypothesis, we used annexin-V staining as a marker to detect early stage of apoptosis after being cultured for 14 hours with or without PFNA (600 μM). Annexin-V is an anticoagulant protein that could bind to phosphatidylserine on the outer leaflet of the cell membrane when apoptosis initiates. Our results showed that the signal of annexin-V within zona pellucida in PFNA-exposed oocytes was much stronger than that in the control oocytes (Fig. 6A). Statistical analysis of the relative intensity of the annexin-V signal showed that the signal level in the PFNA-exposed oocytes was significantly higher than that in the control (2.36 ± 0.19 in the 600 μM PFNA group versus 1.00 ± 0.07 in the control, P < 0.001; Fig. 6B), indicating that PFNA increased early-stage apoptosis level in the PFNA-treated oocytes.

**Figure 6.**
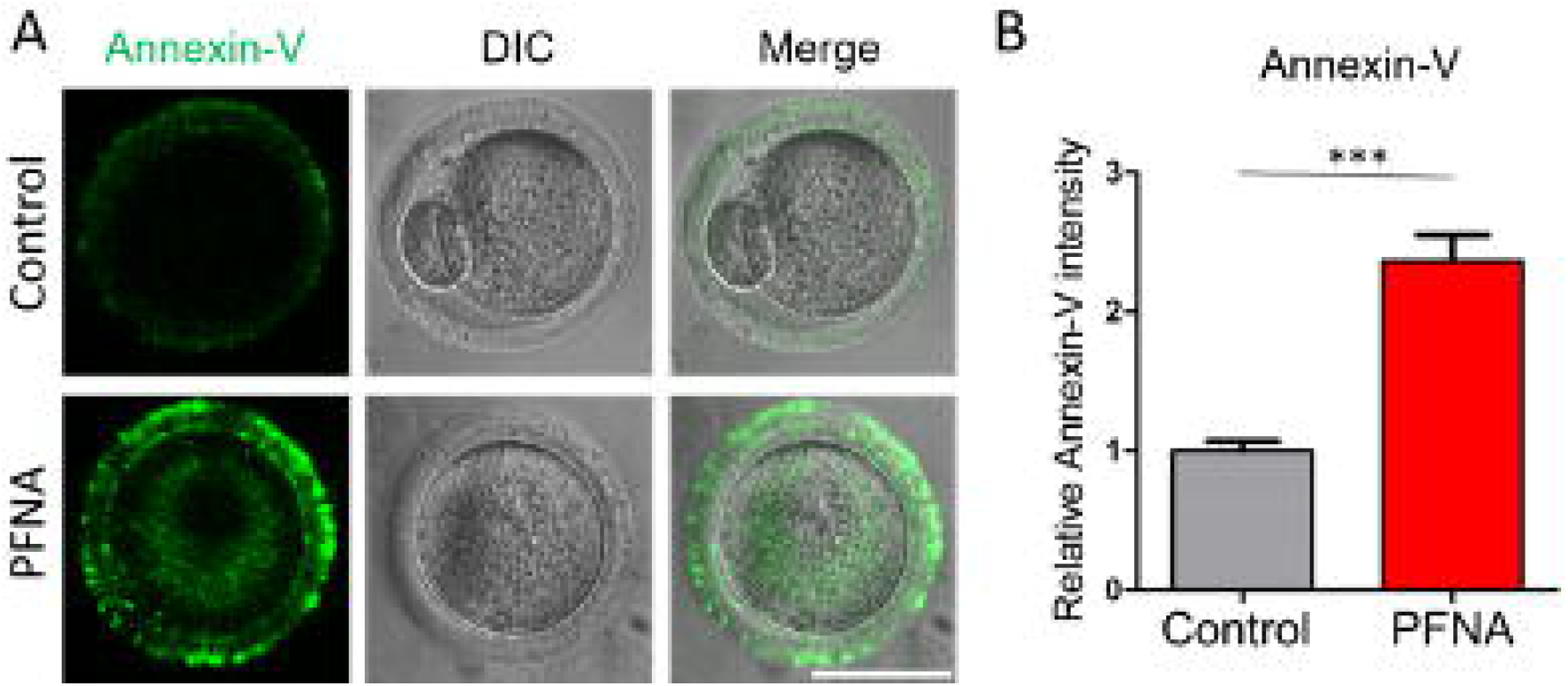
PFNA induced oocyte apoptosis. (A) Representative images showed Annexin-V signal (green) in the control and 600 μM PFNA-treated oocytes. DIC, differential interference contrast. Scale bar, 50 μm. Note: almost all the oocytes in the control group progressed into PBE; while majority of the oocytes in the 600 μl PFNA-treated group arrested in an earlier stage. (B) Quantitative analysis of relative Annexin-V intensity in the control and 600 μM PFNA-exposed oocytes. A total of 42 oocytes in the control and 49 oocytes in the PFNA-treated group were used for Annexin-V signal intensity analysis. Data were presented as mean percentage (mean ± SEM) of at least three independent experiments. t-test, ***P < 0.001, compared with control.

## 4. Discussion

PFNA, a long-chain PFAA, has attracted more and more attention due to its increase in human blood levels and adverse toxicities [12,13]. Recent studies indicated that PFNA exposure inhibited spermatogenesis and lowered sperm quality in male mice [22,23], raising the possibility that PFNA may be a reproductive toxicant. However, whether PFNA exposure causes female reproductive toxicity, especially oocyte maturation failure, was unknown. In this study, we evaluated the toxic effects of PFNA exposure on mouse oocyte maturation *in vitro*. Our results suggest that PFNA impaired the progression of GVBD and inhibited polar-body extrusion (PBE). We further showed that PFNA induced abnormal spindle assembly, chromosome misalignment, elevated ROS levels, and DNA damage.

GVBD allows oocytes to resume their meiotic progress from dictyate arrest. Defects in the progression of GVBD result in a delay or block of meiotic resumption, finally leading to polar body extrusion problems [34,35]. Thus, correct GVBD completion is required for generating mature oocytes. Our results showed that PFNA inhibited GVBD progression in mouse oocytes, indicated by a reduced GVBD rate in the treated oocytes. Notably, the PBE rate was further reduced in the treated oocytes that completed GVBD, indicating that PFNA also inhibited the metaphase I-to-II transition.

To determine the underlying mechanisms of how PFNA disrupted polar-body extrusion in treated oocytes, we checked MI spindles and chromosome alignment because abnormal spindle assembly and chromosome alignment usually contribute to PBE failure in oocytes [36]. Our results showed that PFNA exposure resulted in severe spindle abnormalities as evidenced by disorganized spindles and unclustered MTOCs. Chromosome misalignment was also observed in treated oocytes. In summary, our data indicated that PFNA could impede PBE by disrupting spindle formation and chromosome alignment.

Maturing oocytes have more mitochondria compared to somatic cells [37]. Mitochondria are known to be vulnerable to and targeted by environmental pollutants [38]. Xin et al. [18] showed that PFNA exposure resulted in hyperpolarization of mitochondria in lung cells. Similarly, our results showed that PFNA elevated mitochondrial membrane potential in oocytes. In addition, mitochondrial dysfunction was indicated by abnormal mitochondrial distribution in the PFNA-exposed oocytes. Therefore, we concluded that the mitochondria are the targeted organelles of PFNA in oocytes and that PFNA induces mitochondrial dysfunction in many cell types.

Reactive oxygen species (ROS) are mainly generated in mitochondria during energy production. Mitochondrial dysfunction and increased membrane potential usually elevate ROS levels and lead to oxidative stress in cells [39]. Oxidative stress can affect meiotic progression as well as induce maturation failure in oocytes [40–42]. Our results showed that PFNA significantly elevated the ROS levels and caused DNA damage via oxidative stress. This strongly suggests that PFNA impedes oocyte maturation by inducing oxidative stress, which is consistent with a previous PFNA toxicological study in lung cells [18]. In addition, DNA damage in mouse oocytes can trigger MI arrest causing PBE failure [43,44]. Therefore, we concluded that PFNA-induced DNA damage is another cause of maturation failure in exposed oocytes. Although we demonstrated that oxidative stress induced by PFNA exposure could contribute to maturation defects in oocytes, other factors controlled by mitochondria, such as ATP generation and Ca^2+^ homeostasis, could also govern meiotic progression in oocytes. Therefore, further studies are needed to clarify the relationship between mitochondrial dysfunction and the inhibition of meiotic progression in PFNA-treated oocytes.

Metaphase II (MII) spindle assembly is one of the indicators of the quality of matured oocytes; abnormal MII spindle assembly is highly related with poor pregnancy outcomes in mammals [45]. Our results showed that PFNA resulted in severe spindle abnormalities and chromosome misalignment in exposed MII oocytes. Therefore, we conclude that PFNA exposure also affected the quality of matured oocytes by disrupting MII spindle formation. Indeed, the exposure of PFNA and perfluorodecanoic acid (PFDA) is associated with miscarriage of pregnant women [46]. Higher PFNA is also positively correlated with longer time to pregnancy [47]. However, no associations with PFOA and PFOS were found in these two studies [46,47].

We found that the environmentally relevant dose (3 nM) [11] of PFNA does not affect mouse oocyte maturation *in vitro*. However, the exposure period (< 15 hours) of this study is much shorter than the real exposure scenarios - the half-lives of PFAA in human serum last for several years [48]. Therefore, the impacts of PFNA are significant for long-lived cells, especially for oocytes that can sit in ovaries for decades. PFNA is found in human follicular fluid and bioaccumulative [49], which suggests that the PFNA levels in the oocytes might have a big range. Clinical studies also indicated that PFNA exposure was associated with pregnancy defects [46,47]. Here, we explored a concentration range of PFNA (3 nM – 600 μM) [18] to investigate the mechanisms of oocyte cytotoxicity. Thus, this study can help us better understand PFNA toxicity on oocyte maturation and PFNA-induced reproductive problems.

## 5. Conclusions and Future Directions

In conclusion, our results showed that PFNA impeded mouse oocyte maturation via inhibiting GVBD, disrupting spindle assembly, causing mitochondrial dysfunction, oxidative stress, DNA damage, and early stage of apoptosis. Our study provides new insights into the toxic effects of PFNA exposure on mouse oocyte maturation *in vitro*. However, in real scenario, oocytes from humans and animals are exposed to much lower levels of PFNA for years, compared to the dose (600 μM) we used in this study, which cannot be studied *in vitro*. Thus, *in vivo* studies, especially with a low PFNA dose and a long treatment period, are still needed to further explore the impacts of PFNA exposure on oogenesis.

## Acknowledgement

We would like to thank Drs. Indrani C Bagchi and Gee Lau for kindly sharing reagents. We want to thank the laboratories of Drs. Michael J Spinella and Sarah Freemantle for their technical support and Dr. Jodi Flaws for her comments on this manuscript. This work was supported by National Institutions of Health (NIH) R00 HD082375 and NIH R01 GM135549.

## Author contributions

**XJ and NL:** Conceptualization, Methodology, Validation, Investigation, Formal analysis, Writing-Original Draft. **YX:** Investigation, Writing-Review & Editing. **HQ:** Conceptualization, Resources, Project administration, Supervision, Writing-Review & Editing, Funding acquisition.

