## Supplemental Figure 1 for "Perfluorononanoic acid impedes mouse oocyte maturation by inducing mitochondrial dysfunction and oxidative stress"

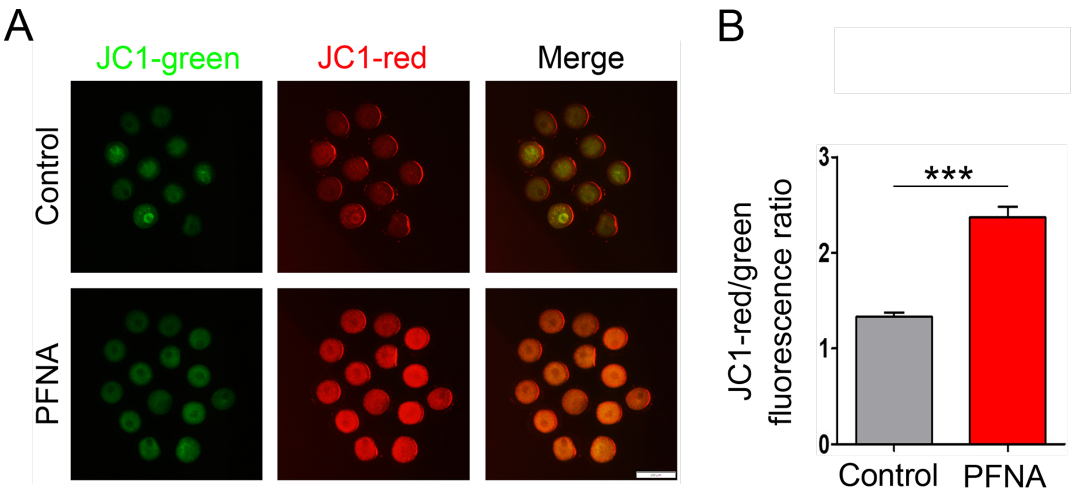


**Supplemental Figure 1. PFNA exposure elevated mitochondrial membrane potential.** (A) Staining of JC-1 (an indicator of mitochondrial membrane potential) in the control and 600 μM PFNA-treated oocytes. (B) The ratio of fluorescence intensity of JC1-red/green in the control and 600 μM PFNA-exposed oocytes. A total of 33 oocytes in the control and 41 oocytes in the 600 μM PFNA-treated groups were analyzed for JC-1 analysis. Data were presented as mean ± SEM of at least three independent experiments. t-test, $^{**}P < 0.01$ and $^{***}P < 0.001$, compared with control.
